# Prevalence and antibiotic resistance of *Salmonella* spp. in South Punjab-Pakistan

**DOI:** 10.1101/2020.04.15.042705

**Authors:** Aftab Qamar, Tariq Ismail, Saeed Akhtar

**Affiliations:** Institute of Food Science and Nutrition Bahauddin Zakariya University Multan

**Keywords:** Antibiotic Resistance, *Salmonella*, Raw milk, Pakistan

## Abstract

Present study aimed at investigating the magnitude of the prevalence and antibiotic resistance among four *Salmonella* spp. i.e. *S. typhi*, *S. paratyphi* A, *S. paratyphi* B and *S. typhimurium,* present in raw milk and environment samples from five districts of southern part of the Province of Punjab in Pakistan i.e. Multan, Bahawalpur, Lodhran, Dera Ghazi Khan and Muzaffargarh. A total of 3000 raw milk and environment samples were collected from these sites and analyzed for detection and confirmation of *Salmonella* spp. Extent of antibiotic resistance was also determined and classified as resistant, intermediate and susceptible. *S. typhimurium* was found to be more prevalent in Multan region as compared to other districts. Increased emergence of antibacterial resistance among *Salmonella* spp. from raw milk samples was noted. Amongst 13 tested antibiotics, chloramphenicol and ofloxacin were found to be the most susceptible against *Salmonella* spp. Present study suggested *serious* interventions by relevant government agencies to ensure best practices in animal husbandry at farm level and sagacious use of antibiotics for the containment of antimicrobial resistance in *Salmonella* spp.

## Introduction

Traditionally, raw milk has been considered as a source of nutrients by a variety of farming families and workers which they believe, are destroyed during any kind of processing [1] however raw milk and milk products are reported to be the best breeding sites for pathogenic microbial strains hence safety of these products turns up as a challenge given the direct contact of raw milk with contaminated surfaces such as air, soil, workers hygiene, feces, grass and excretion from the udder of an infected animal and milking equipments in production and processing areas [1].

*Salmonella* has been considered as the major foodborne pathogen leading to an upsurge in enteric infection cases. Three groups of *Salmonella* serotypes have been considered responsible for causing distinctive clinical syndromes including typhoid fever, enteritis and bacteremia [2]. Likewise, infections by non-typhoid *Salmonella* serovars have been shown to result in acute gastroenteritis with extra intestinal localized infections that may eventually affect some organs [3]. Reportedly, 99% *Salmonella* infections in humans are associated with strains in the O-antigen serogroups such as serogroups A, B, C1, C2, D and E of *S. enterica* subspecies entericae [4]. Mechanistically, the onset of disease proceeds with intestinal phase once the food contaminated with typhoid and *Salmonella enteritis* is ingested [5].

Recently, emergence of multidrug-resistant (MDR) *S. enterica* serovars, including resistance to quinolone group (fluoroquinolones) and the third generation antibiotics (cephalosporin) has led to serious public health issues throughout the world. One study demonstrated emergence of antimicrobial resistance among *Salmonella* during animal farming, indicating 10% of the isolates being resistant against one or more tested antimicrobials commonly used to treat *Salmonella* infections e.g. cephalosporin and fluoroquinolones[6]. Sufficient evidence is available to support the emergence of antibiotic resistance among *Salmonella* strains which is directly associated with intensive use of antibiotics to treat *Salmonella* infections and incorporation of growth promoters in animal feed [7,8]. The wider distribution of multidrug resistant (MDR) *Salmonella* spp. in foods have been reported by various researchers worldwide [9–11]. A substantial body of literature confirmed the prevalence of antibiotic resistance among foodborne pathogens especially in milk and milk products and such a resistance pattern also came out in *Salmonella* spp. in last few decades as a major public health issue worldwide [12,13].

*S. enterica* serovars *typhi* and *paratyphi* (A, B and C) have been reportedly developing resistance against a range of antibiotics thereby distressing 21 million people worldwide. Morbidity and mortality rate associated with these microbial infections had been much higher on account of infections by Salmonella spp. For example more than 14 million cases of enteric fever are reported annually resulting in 135,000 deaths. *S. typhi* and *S. paratyphi* infections have been shown to be more prevalent in South Asian regions where excessive use of antibiotic has been exacerbating the emergence of multidrug resistance in these strains [14,15]. Multi-drug resistance (MDR) *Salmonella* serotypes have become widespread in developed economies as well including USA. Treatment cost of infections from antibiotic resistant bacteria amounts to 4-5 billion US dollars annually. In addition to substantial financial losses caused in disease management, antibiotic resistant pathogens have been hampering international trade owing to threats of cross borders proliferation of infectious diseases [16].

Available evidence suggest increased prevalence of *Salmonella* spp. in foods especially raw milk and milk products leading to a surge in the onset of infections among humans and the farm animals. Resultantly, the injudicious and indiscreet use of antibiotics to treat such infections has been engendering heightened multidrug resistance among bacterial strains. Besides that, no epidemiological surveillance, monitoring and control of pathogenic microbes and associated microbiological infections is in place. The objective of the present study was to scale the prevalence of *Salmonella* spp. at farms in South Punjab and to ascertain the extent of development of antibiotic resistance in *Salmonella* spp. isolated from raw milk and farm environment. Aftermaths of this study would provide a baseline information for the policy makers, farm owners, dairy industry and stake holders to initiate plans for mitigating microbiological food safety issues and corresponding disease burden in the region.

## Materials and Methods

A cross-sectional study was designed to find out the prevalence of *Salmonella* spp. in raw milk and environmental samples collected from five major districts of South Punjab (Fig.1). A total of 3000 samples were collected from towns/tehsils of these districts and detection and isolation of *Salmonella* spp. were carried out to estimate the extent of the prevalence of *Salmonella* spp. i.e. *S. typhi, S. paratyphi A, S. paratyphi B and S. typhimurium*. The isolated *Salmonella* strains were also tested against antibiotics i.e. Gentamicin (GEN), Chloramphenicol (CPL), Ampicillin (AMP), Oxytetracycline (OTC), Ciprofloxacin (CIP), Ofloxacin (OFL), Amoxicillin (AMX), Cefuroxime (CXM), Ceftazidime (CZA), Cefepime (CPE), Imipenem (IMP), Trimethoprim (TMP), Moxalactam (MOX), to figure out the resistance pattern in the organisms. The selection of tested antibiotics was made, based on the present therapeutic use of these antibiotics to treat *Salmonella* infections in humans and farm animals.

**Figure-1.**
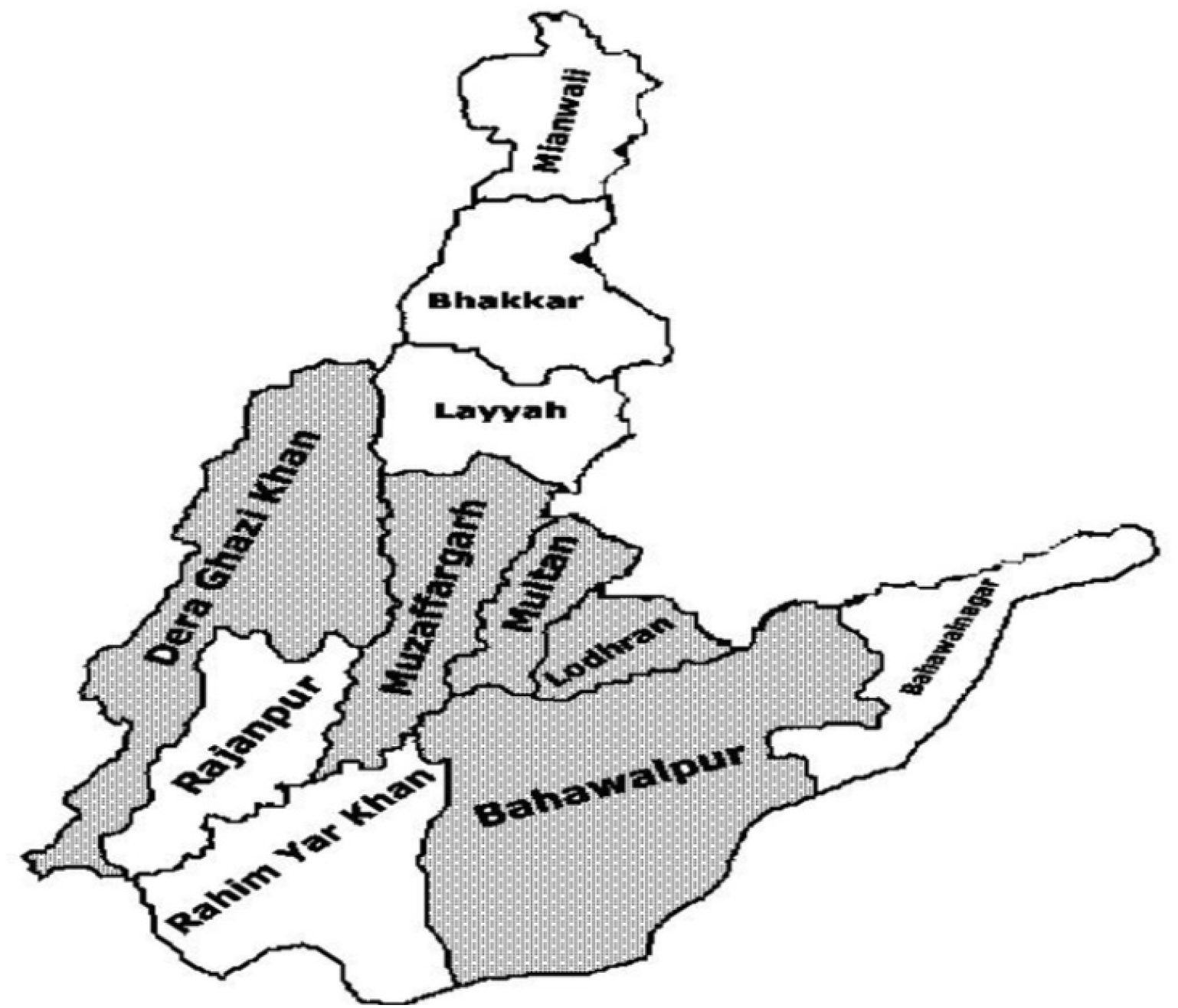
Map showing five major districts (shaded) of South Punjab – Pakistan covered for surveillance and antibacterial resistance in Salmonella isolates from raw milk and environment;

A total of 15 raw milk and 135 environment samples (manure, soil, feed, bedding, potable water, milk container, personnel and animals’ teat) were collected from dairy farms, bicycle milk vendors and raw milk outlets from peri-urban areas of each targeted site. Raw milk (approx. 50 ml in sterilized dry bottles) and environmental samples (almost 100 g in sterilized zip lock bags) were collected, tightly sealed, kept in ice box and immediately shipped to laboratory for analyses. Procedure from ISO 6579:2002 standard (Microbiology of food and animal feeding stuffs) guidelines were followed for detection, isolation and confirmation of *Salmonella.*

All *Salmonella* positive samples were examined to measure the gravity of antibiotic resistance using Hardy DiskTM Antimicrobial Sensitivity Test (AST) (Hardy Diagnostic, Santa Maria, CA). Isolates were evaluated for resistance to a panel of thirteen antibiotics by measuring zones of inhibition with a meter ruler. This test was performed twice to calculate the means for inhibition zone and isolates were declared resistant, intermediate and susceptible against the employed antibiotics according to the Clinical & Laboratory Standard Institute (CLSI) guidelines. All chemicals and bacterial culture media were of analytical-reagent grade if otherwise noted.

The data for prevalence of *Salmonella* spp. so obtained were subjected to statistical analysis and positive and negative samples were taken to calculate percentage prevalence of different *Salmonella* spp. in these samples. The prevalence was tested with Chi squared test at 5 % level of significance.

## Results

Data presented in Table 1 depict % prevalence of *S typhi* (8.33%)*, S. paratyphi* A (2.78%), *S. paratyphi* B (3.67%) and *S. typhimurium* (10.89%) isolated from 233 positive sample screened from 900 raw milk and environmental samples from six towns of Multan district i.e. *Band Bosan, Shah Rukn Alam, Musa Pak Shaheed, Sher Shah, Shuja Abad and Jalal Pur Pir Wala*. Comparably, *S. tyhpimurium* remained to be the most frequent *Salmonella* serovar, however variability in prevalence rate was non-significant (*P>0.05*). All six towns significantly differed (*P>0.05*) for prevalence rate with highest prevalence of *S. tyhpimurium* (16.67 %) in *Shuja Abad.* This site indicated overall highest prevalence (39.33%) of *Salmonella* spp. with higher number (*n=59*) of positive samples followed by *Band Bosan* town with29.33% (*n=43*) and *Sher Shah* with 26.67% (*n=40*). Relative to these sites, *Shah Rukn Alam* was identified as microbiologically safe area with 14.67% (*n=22*) prevalence of *Salmonella* spp. (Table 1).

**Table 1.** Prevalence (%) of *Salmonella* spp. isolated from raw milk and environment samples in dairy farms from district Multan

| Towns | TS | PS | *S. typhi*<br>n (%) | *S. paratyphi A*<br>n (%) | *S. paratyphi B*<br>n (%) | *S. typhimurium*<br>n (%) | Total<br>Prevalence<br>(%) |
| --- | --- | --- | --- | --- | --- | --- | --- |
| **BB** | 150 | 43 | 12 (8.0) | 3 (2.0) | 8 (5.33) | 20(13.33) | **29.33** |
| **SRA** | 150 | 22 | 7 (4.67) | 4 (2.67) | 2 (1.33) | 9 (6.0) | **14.67** |
| **MPS** | 150 | 31 | 15 (10.0) | 4 (2.67) | 3 (2.0) | 9 (6.0) | **20.67** |
| **SS** | 150 | 40 | 11 (7.33) | 5 (3.33) | 7 (4.67) | 17 (11.33) | **26.67** |
| **SAB** | 150 | 59 | 19 (12.67) | 6 (4.0) | 7 (4.67) | 25 (16.67) | **39.33** |
| **JPPW** | 150 | 38 | 11(7.33) | 3 (2.0) | 6 (4.0) | 18 (12.0) | **25.33** |
| **Total** | **900** | **233** | **75 (8.33)** | **25 (2.78)** | **33 (3.67)** | **98 (10.89)** | **25.89** |

Town ; BB: Band Bosan, SRA: Shah Rukne Alam, MPS; Musa Pak Shaheed, SS: Sher Shah, SAB: Shuja Abad, JPPW: Jalal Pur Pir Wala TS; Total number of samples, PS; Total number of positive sample, n = Number of positive samples of respective spp

Amidst five experimental sites in district Bahawalpur, *Ahmad Pur East* was shown to elicit the highest prevalence (%) for all four *Salmonella* spp. with *S*. *typhi* being more visible (Table 2). *S. typhi* also turned up as the most prevalent *Salmonella* spp. i.e. 11.8% (*n=89*) in Bahawalpur district as a whole, followed by S. *typhimurium* i.e. 10.0 % (*n=75*). Notwithstanding, prevalence (%) of *S. paratyphi* A&B marked a non-significant difference (*P>0.05*) and they appeared to be the least prevalent *Salmonella* spp. i.e. 3.2 % (*n=24*) and 2.8% (*n=21*) respectively in the region. When it came to the town level prevalence rate, *Hasil Pur* seemed to have been microbiologically the least tainted site in district Bahawalpur. A total of 209 samples (27.87%) were found positive for four strains of *Salmonella* isolated from raw milk and environment (Table 2.)

**Table 2.** Prevalence (%) of *Salmonella* spp. isolated from raw milk and environment samples in dairy farms from district Bahawalpur

| Towns | TS | PS | *S. typhi*<br>n (%) | *S. paratyphi A*<br>n (%) | *S. paratyphi B*<br>n (%) | *S. typhimurium*<br>n (%) | Total Prevalence<br>(%) |
| --- | --- | --- | --- | --- | --- | --- | --- |
| **BWP** | 150 | 37 | 17 (11.33) | 5 (3.33) | 4 (2.67) | 11 (7.3) | **24.67** |
| **APE** | 150 | 56 | 24 (16.0) | 7 (4.67) | 8 (5.33) | 17 (11.33) | **37.33** |
| **HPR** | 150 | 29 | 12 (8.0) | 2 (1.33) | 3 (2.0) | 12 (8.0) | **19.33** |
| **YZN** | 150 | 41 | 13 (8.67) | 6 (4.0) | 3 (2.0) | 19 (12.67) | **27.33** |
| **KPT** | 150 | 46 | 23 (15.33) | 4 (2.67) | 3 (2.0) | 16 (10.67) | **30.67** |
| **Total** | **750** | **209** | **89 (11.8%)** | **24 (3.2%)** | **21 (2.8%)** | **75 (10.0%)** | **27.8** |

Towns ; BWP=Bahawalpur, APE=Ahmad Pur East, HP; Hasil Pur, YZN; Yazman, KPT; Khairpur Tamewali TS; Total number of samples, PS; Total number of positive sample, n = Number of positive samples of respective spp.

Lodhran is relatively a smaller district of South Punjab and is located on northern side of the River Sutlej with its three towns viz *Lodhran, Dunya Pur, Kehror Pakka*. A total of 209 (20.22%) samples from raw milk and environment appeared as positive for *Salmonella* (Table 3). Considering the extent of *Salmonella* contamination in different towns, comparative results for prevalence (%) of *Salmonella* spp. revealed that the environment of sites from district *Dunya Pur*, was the most polluted with *S. typhi* (11.33 %) and *S. typhimurium* (9.33%) both being more prevalent as compared to other *Salmonella* spp. (Table-3).

**Table 3.** Prevalence (%) of *Salmonella* spp. isolated from raw milk and environment samples in dairy farms from district Lodhran

| Towns | TS | PS | *S. typhi*<br>n (%) | *S. paratyphi A*<br>n (%) | *S. paratyphi B*<br>n (%) | *S. typhimurium*<br>n (%) | Total Prevalence<br>(%) |
| --- | --- | --- | --- | --- | --- | --- | --- |
| **LDN** | 150 | 28 | 11 (07.33) | 1 (0.67) | 4 (02.67) | 12 (08.00) | **18.67** |
| **DPR** | 150 | 40 | 17 (11.33) | 5 (3.33) | 4 (02.67) | 14 (09.33) | **26.67** |
| **KPA** | 150 | 23 | 07 (04.67) | 3 (2.00) | 2 (01.33) | 11 (07.33) | **15.33** |
| **Total** | **450** | **209** | **35 (07.78)** | **09 (2.00)** | **10 (02.22)** | **37 (08.22)** | **20.22** |

Towns; LDN; Lodhran, DPR; Dunya Pur, KPA; Kehror Pakka
TS; Total number of samples, PS; Total number of positive sample, n = Number of positive samples of respective spp

Similarly, data presented in Table 4 represent prevalence (%) of *Salmonella* spp. from *Dera Ghazi khan* and *Taunsa Sharif* suggesting *S*. *typhi* to be the most prevalent (11.0%) strain followed by *S. typhimurium* (10.33%). Town wise total percentage of *Salmonella* spp. in *D.G. Khan* town was 27.33% (*n=41*) whereas *Taunsa Sharif* showed higher rate i.e. 37.33% (*n=56*) with an overall percentage of 32.33% in the whole district. A total of 97 (32.3%) out of 300 raw milk and environment samples were positive for *Salmonella* spp. in *D.G. Khan*. Differences among *Salmonella* spp. were non-significant (*P>0.05*) with regard to prevalence in towns of D.G. Khan region as shown in Table 4.

**Table 4.** Prevalence (%) of *Salmonella* spp. isolated from raw milk and environment samples in dairy farms from district Dera Ghazi Khan

| Towns | TS | PS | *S. typhi*<br>n (%) | *S. paratyphi A*<br>n (%) | *S. paratyphi B*<br>n (%) | *S. typhimurium*<br>n (%) | Total Prevalence<br>(%) |
| --- | --- | --- | --- | --- | --- | --- | --- |
| **DGK** | 150 | 41 | 12 (08.00) | 05 (03.33) | 07 (04.67) | 17 (11.33) | **27.33** |
| **TSA** | 150 | 56 | 21 (14.00) | 06 (04.00) | 15 (10.00) | 14 (09.33) | **37.33** |
| **Total** | **300** | **97** | **33 (11.00)** | **11 (03.67)** | **22 (07.33)** | **31 (10.33)** | **32.33** |

Towns; DGK: D.G. Khan, TSA: Taunsa Sharif
TS; Total number of samples, PS; Total number of positive sample, n = Number of positive samples of respective spp

Muzaffargarh is one the known districts in Dera Ghazi Khan division of Punjab in Pakistan. Muzaffargarh city is located on the banks of the Chenab River. Out of 600 samples screened for *Salmonella*, 31.33% (*n=188*) samples were found positive with the most prevalent *Salmonella* spp. *S. typhimurium* 13.67% (*n=82*) followed by *S. typhi* 7.83% (*n=47*), *S. paratyphi* B 6.0% (*n=36*) while *S. paratyphi* A accounted for 3.83% (*n=23*) being the least prevalent *Salmonella* spp. (Table 5). Amongst all experimental sites, *Ali Pur* was observed to be highly infected with 40.67% (*n=61*) prevalence of *Salmonella* spp. followed by *Kot Addu* 32.67% (*n=49*), Jatoi 28.67% (*n=43*) and *Muzaffargarh* 23.33% (*n=35*). *Kot Addu* and *Ali Pur* showed high prevalence rate of *S. typhi* and *S. typhimurium* with 10.0% (*n=15*) and 18.0% (*n=27*), respectively. Similar prevalence rate of *S. paratyphi* A 3.3% (*n=5*) was observed in *Muzaffargarh* and *Kot Addu* areas whereas least occurrence (2.7%) of *S. paratyphi* B was recorded in *Muzaffargarh* town. The results presented in Table 5 showed differences among *Salmonella* spp. as non-significant (*P>0.05*).

**Table 5.** Prevalence (%) of *Salmonella* spp. isolated from raw milk and environment samples in dairy farms from district Muzaffargarh

| Towns | TS | PS | *S. typhi*<br>n (%) | *S. paratyphi A*<br>n (%) | *S. paratyphi B*<br>n (%) | *S. typhimurium*<br>n (%) | Total Prevalence<br>(%) |
| --- | --- | --- | --- | --- | --- | --- | --- |
| **MZG** | 150 | 35 | 10 (06.67) | 5 (3.33) | 4 (2.67) | 16 (10.67) | **23.33** |
| **KAU** | 150 | 49 | 15 (10.00) | 5 (3.33) | 7 (4.67) | 22 (14.67) | **32.67** |
| **APR** | 150 | 61 | 12 (08.00) | 7 (4.67) | 15 (10.0) | 27 (18.00) | **40.67** |
| **JTI** | 150 | 43 | 10 (06.67) | 6 (04.00) | 10 (6.67) | 17 (11.33) | **28.67** |
| **Total** | **600** | **188** | **47 (07.83)** | **23 (03.83)** | **36 (06.00)** | **82 (13.67)** | **31.33** |

Towns; MGR=Muzaffargarh, KAU; Kot Addu, JTI; Jatoi, APR; Ali Pur
TS; Total number of samples, PS; Total number of positive sample, n = Number of positive samples of respective spp

## Discussion

### Prevalence of *Salmonella* spp

The area under study has been one of the most distressed regions of Pakistan in terms of health care system and provision of medical facilities. Poverty remains to be a challenge which results in increased disease burden. Present study reflected higher rate of prevalence of *Salmonella* spp. in South Punjab indicating heighted incidences of salmonellosis. For example, overall prevalence of *Salmonella spp*. in all towns of Multan district was noted to be 25.89% (Table 1) and similar findings were also presented byRahman *et al*. [17] who reported 21.89% *Salmonella* spp. in different samples. Other studies demonstrated the prevalence levels of *Salmonella* to be ranging from 7.61% to 11.9 % attributing the same to a variety of factors important being hygiene, sanitation and training of the food handling staff [18,19]. Variability and significant differences in temperature at experimental sites in the present study could be a key determinant for difference in level of prevalence of *Salmonella* spp.

Our data revealed prevalence of *S. typhi* isolated from raw milk samples in district Bahawalpur to be to the tune of 11.9% (Table 2). Similar results were presented by Addis *et al.* [20] who reported *Salmonella* at 10.76% (*n*=21/195) either from milk or feces samples. Similarly, 35.71% milk samples were found to be positive for *S. typhi* in Bangladesh [21]. Apart from Southern Punjab regions, more reports are available to signify the overwhelming effects of *S. enteritidis* among a number of population groups. Aikin to other districts, *S. typhi* and *S. typhimurium* indicated the similar trend for prevalence irrespective of the sampling sites and sample type in district Lodhran which is a proxy of overall environment at dairy farms in the area (Table 3)

Current study further revealed 32.3% *Salmonella* samples being positive in district D.G. Khan. The results of present study are in agreement with the finding of Pangloli *et al.* [22] who isolated 40-92% *Salmonella* spp. from animal and environment samples. High prevalence of *Salmonella* spp. was ascribed to the poor hygienic condition of dairy farms, seasonal variation and improper personnel cleanliness. Our results further confirmed prevalence of *S. typhi* (11.0%) in D.G. Khan being less than extent of prevalence reported by Soomro *et al.* [23] who identified high prevalence of *Salmonella enteritidis* from chicken meat samples. The low prevalence of *Salmonella* spp. in this area was of *S. paratyphi* A with a prevalence rate of 3.67%. Almost identical results were obtained by other researchers who isolated *Salmonella* from Kariesh cheese samples [24]. Increased prevalence of *Salmonella* spp. was also reported by Ghada *et al.* [25]and Wallaa, [26] who observed isolated *Salmonella* spp. from milk and cheese at 10% and 4% respectively.

Comparing the town wise prevalence of *Salmonella* in Muzaffargarh, Ali *pur* was shown to indicate higher positive samples of *Salmonella (*Table 5). Prevalence rate of *Salmonella* however might not be attributed to any specific determinant and no relationship with respect to prevalence rate and region was established except the reasons described above i.e. hygiene, sanitation, animals’ health and training of the staff. Data are not scant to indicate that the prevalence of *Salmonella* at farms is not farm type specific e.g. beef cattle farm or dairy farms. These researchers were of the view that variation in prevalence might be a result of location of the farms and the focus on pathogen isolation from fecal or other animal-based samples [27–30]

Our results have further substantiated that the difference in prevalence of *Salmonella* spp. and the sources statistically differed with variability in region and source type however, a striking difference in prevalence rate of *Salmonella* spp. among regions could not be established to ascertain a baseline for *Salmonella* occurrence in selected districts. Overall results of this study demonstrated that no raw milk and environmental sample from selected sites might be considered up to the defined standards with respect to microbiological safety of the food and control and monitoring of dairy farms. Murinda *et al.* [29] reported 2.2% of bulk tank milk samples contaminated with *Salmonella*, attributing the presence of *Salmonella* in tanks to be the result of cross-contamination from milking environmental sites instead of animal sites. A few recent studies with small sample size indicated *Salmonella* to be present in raw farm bulk milk at 12% [31]. Results from a similar recent study from Ghana explicated reduced prevalence of *Salmonella enterica* in cow milk i.e. 7.3% [32].

A perusal of earlier studies to contemplate and compare the extent of prevalence of *Salmonella* in South Asian regions portrayed that the prevalence rate in dairy and dairy products was more or less the same. Findings from Singh *et al.* [33]and Pant *et al.* [34] substantiated a kind of similar prevalence rate in India. Kaushik *et al.* [35] observed similar prevalence rate of *Salmonella* in market milk samples in Patna, Bihar. Bangladesh as a region in subcontinent was not an exception for higher *Salmonella* prevalence where the presence of *S. typhi* was found to be 35.17% in vendor’s milk. More studies confirmed these results showing *Salmonella* prevalence to the tune of 9.5% and 4.2% [21,36,37] This variation justified high prevalence of *Salmonella* in various South Asian regions especially those located in subcontinent i.e. Pakistan and India because cultural, atmospheric and social conditions were quite the same therefore we might have witnessed the prevalence level being reported from these areas to be more or less similar.

### Antimicrobial Resistance in *Salmonella* spp

Data presented in Table 6 revealed the extent of resistance of *Salmonella* spp. against an array of antibiotics. *S typhi* emerged as a highly resistant *Salmonella* strain against OTC (70.11%) followed by AMP (38.79%), TMP (33.45%), CPl (29.54%) and AMX (28.11%) whilst same strain had shown to be the least resistant against OFL (0.00%), MOX (0.00%) and CPE (1.07%) suggesting these antibiotics to be employed against *S. typhi* infections. Four antibiotics viz OFL, CXM, IMP and MOX were noted to be remarkably effective against *S. paratyphi A* infection whereas this strain was identified to be highly resistant against OTC (25.84%) and TMP (47.19%). *S. paratyphi A* had also shown increased tendency towards switching over from sensitivity zone to intermediate level resistance against OTC (24.72%), TMP (21.35%), CXM (19.10%), and MOX (14.61%) suggesting a more cautious use of these antimicrobials against *S paratyphi* A infection.

**Table 6.** Antimicrobial susceptibility pattern of *Salmonella* isolates from raw milk and environment samples in dairy farms.from South Punjab-Pakistan

| Antibiotic<br>(µg) | *S. typhi* n (%) |  |  | *S. paratyphi A* n (%) |  |  | *S. paratyphi B* n (%) |  |  | *S. typhimurium* n (%) |  |  |
| --- | --- | --- | --- | --- | --- | --- | --- | --- | --- | --- | --- | --- |
|  | Sen | Int | Res | Sen | Int | Res | Sen | Int | Res | Sen | Int | Res |
| GEN (10) | 228<br>(81.14) | 34<br>(12.10) | 19<br>(6.76) | 78<br>(87.64) | 7<br>(7.87) | 4<br>(4.49) | 114<br>(94.21) | 7<br>(5.79) | 0<br>(0.00) | 267<br>(81.40) | 28<br>(8.54) | 33<br>(10.06) |
| CPL (30) | 152<br>(54.09) | 46<br>(16.37) | 83<br>(29.54) | 64<br>(71.91) | 19<br>(21.35) | 6<br>(6.74) | 84<br>(69.42) | 22<br>(18.18) | 15<br>(12.40) | 224<br>(68.29) | 73<br>(22.26) | 31<br>(9.45) |
| AMP (10) | 123<br>(43.77) | 49<br>(17.44) | 109<br>(38.79) | 62<br>(69.66) | 8<br>(8.99) | 19<br>(21.35) | 95<br>(78.51) | 15<br>(12.40) | 11<br>(9.09) | 175<br>(53.35) | 66<br>(20.12) | 87<br>(26.52) |
| OTC (30) | 52<br>(18.51) | 32<br>(11.39) | 197<br>(70.11) | 44<br>(49.44) | 22<br>(24.72) | 23<br>(25.84) | 63<br>(52.07) | 26<br>(21.49) | 32<br>(26.45) | 129<br>(39.33) | 86<br>(26.22) | 113<br>(34.45) |
| CIP (05) | 239<br>(85.05) | 11<br>(3.91) | 31<br>(11.03) | 78<br>(87.64) | 2<br>(2.25) | 9<br>(10.11) | 114<br>(94.21) | 5<br>(4.13) | 2<br>(1.65) | 221<br>(67.38) | 77<br>(23.48) | 30<br>(9.15) |
| OFL (05) | 254<br>(90.39) | 27<br>(9.60) | 0<br>(0.00) | 84<br>(94.38) | 5<br>(5.62) | 0<br>(0.00) | 103<br>(85.12) | 18<br>(14.88) | 0<br>(0.00) | 302<br>(92.07) | 26<br>(7.93) | 0 (0.00) |
| AMX(30) | 168<br>(59.79) | 34<br>(12.10) | 79<br>(28.11) | 57<br>(64.04) | 19<br>(21.35) | 13<br>(14.61) | 82<br>(67.77) | 16<br>(13.22) | 23<br>(19.01) | 262<br>(79.88) | 41<br>(12.50) | 25<br>(7.62) |
| CXM (30) | 246<br>(87.54) | 23<br>(8.19) | 12<br>(4.27) | 72<br>(80.90) | 17<br>(19.10) | 0<br>(0.00) | 101<br>(83.47) | 13<br>(10.74) | 7<br>(5.79) | 224<br>(68.29) | 88<br>(26.83) | 16<br>(4.88) |
| CZA (30) | 208<br>(74.02) | 39<br>(13.88) | 34<br>(12.10) | 70<br>(78.65) | 14<br>(15.73) | 5<br>(5.62) | 106<br>(87.60) | 15<br>(12.40) | 0<br>(0.00) | 297<br>(90.55) | 26<br>(7.93) | 0 (0.00) |
| CPE (30) | 252<br>(89.68) | 26<br>(9.25) | 3<br>(1.07) | 70<br>(78.65) | 14<br>(15.73) | 5<br>(5.62) | 93<br>(76.86) | 24<br>(19.38) | 4<br>(3.31) | 273<br>(83.23) | 46<br>(14.02) | 9 (2.74) |
| IMP (10) | 240<br>(85.41) | 28<br>(9.96) | 13<br>(4.63) | 87<br>(97.75) | 2<br>(2.25) | 0<br>(0.00) | 109<br>(90.08) | 12<br>(9.92) | 0<br>(0.00) | 312<br>(95.12) | 16<br>(4.88) | 0 (0.00) |
| TMP (25) | 137<br>(48.75) | 50<br>(17.79) | 94<br>(33.45) | 28<br>(31.46) | 19<br>(21.35) | 42<br>(47.19) | 75<br>(61.98) | 31<br>(25.62) | 15<br>(15.40) | 204<br>(62.20) | 51<br>(15.55) | 73<br>(22.26) |
| MOX (10) | 250<br>(88.97) | 31<br>(11.03) | 0<br>(0.00) | 76<br>(85.39) | 13<br>(14.61) | 0<br>(0.00) | 80<br>(66.12) | 41<br>(33.88) | 0<br>(0.00) | 309<br>(94.21) | 19<br>(5.79) | 0 (0.00) |

GEN; Gentamicin, CPL; Chloramphenicol, AMP; Ampicillin, OTC; Oxytetracycline, CIP; Ciprofloxacin, OFL; Ofloxacin, AMX; Amoxicillin, CXM; Cefuroxime, CZA; Cefazidime, CPE; Cefepime, IMP; Imipenem, TMP; Trimethoprim, MOX; Moxalactam Sen; Sensitive, Int; Intermediate, Res; Resistant Numbers in parenthesis indicate percentage prevalence

Our results indicated that *S. paratyphi A* had not yet acquired multidrug resistance against these antibiotics, therefore these drugs could be equally applied as treatment options against illness caused by this microorganism. *S paratyphi* B has almost manifested similar patterns for antibiotic resistance against the tested antibiotics as that of *S paratyphi* A however, the microbe depicted increased sensitivity against GEN (94.21%) and CZA (87.60%) over *S paratyphi* A (Table 6). Our results further demonstrated that S *paratyphi* B was shown to make a rapid transition from its extant sensitivity to developing resistances against MOX (33.88 %) and TMP (25.62%). Comparing *S. typhimurium* with rest of the three *Salmonella* spp. tested for development of antibiotic resistance against 13 antibiotics as mentioned in materials & method section, this strain had exhibited nearly a similar response with being least resistant against OFL, MOX and IMP in addition to CZA (Table 6). Relying on the data obtained in our study, an overall picture of the scale of emergence of antibiotic resistance among *Salmonella* spp. and efficacy of the 13 antibiotics tested against these microorganisms, we suggest OFL and MOX to be the most promising drugs of this time to treat *Salmonella* infections because most of the other antibiotics were shown to be in a transitional phase and are consistently losing their effectiveness against emerging and re-emerging microbes.

Researchers have recently ascribed the presence of antibiotic residues and antibiotic resistance bacteria in the animals’ manure to be the underlying cause of increased spread of antibiotic resistance. Besides, they reported a rise in antibiotic susceptibility among dairy manure isolates of bacterial pathogens with 15% of tested bacteria to be resistant against some antibiotics [38] Most of the bacterial strains have been undergoing genetic modification for evolving resistance on account of indiscriminate and injudicious use of antibiotics for treating animal and human infections. Results of the present study demonstrate similar tendencies as all five experimental sites were shown to have been contaminated with *Salmonella* spp. A similar study depicted the same picture suggesting *Salmonella* isolates from lactating cows, individuals handling them and the environment to be resistant to at least one of the tested antibiotics with 100% to ampicillin. Ciprofloxacin and amoxicillin appeared to be relatively effective as isolates were sensitive to these drugs [39]. More recently, researchers confirmed *Salmonella enterica* isolates from milk to be increasingly resistant to erythromycin (86.0%). Investigators further recorded susceptibility pattern as ciprofloxacin (100.0%), chloramphenicol (91.0%), ceftriaxone (91.0%), tetracycline (86.0%) and ampicillin (86.0%) attributing the increased emergence of resistance to imprudent and indiscreet exploitation of antimicrobials to treat animals against infectious diseases in dairy farms in Ghana and Uruguay [32,40]. Lately, Sobur *et al.* [41] delineated an upsurge in resistance among *Salmonella* spp. against a number of antibiotics including oxytetracycline, tetracycline, erythromycin, azithromycin, and ertapenem. Researcher corroborated that *Salmonella* spp. were widely distributed in dairy farms and their environment and this scenario called for one health approach to override the growing health risks. They suggested judicious and wise use of antibiotics among dairy cattle for their treatment against salmonellosis.

## Conclusion

Our study validated increased prevalence of *Salmonella* spp. on dairy farms in Southern Punjab, which is well known for livestock production in Pakistan. Primarily, higher prevalence of *Salmonella* spp. in these regions badly contaminate the farm environment leading to the onset of more frequent infections among humans and farm animals through various channels including raw milk and direct contact with contaminated farm entities. Emergence of antibiotic resistance has been globally recognized as an issue of public health significance and myriad containment strategies are underway however absence of new antimicrobials with increased efficacy has come out as a serious issue that warrants grave attention of the global health professionals. Available treatment options remain to be the conventional antibiotics being injudiciously used for treating *Salmonella* infections, leaving the microbes more resistant against them. Apparently, appropriate documentation and surveillance of bacterial infections and outbreaks badly lack in this region resulting in greater health risks and increased disease burden. Precise, pragmatic and comprehensive strategies and initiatives have to be brought forward for the control of *Salmonella* infections and the containment of multi drug resistance at regional and national level.

## Acknowledgement

Authors are thankful to Higher Education Commission of Pakistan for supporting research of Mr. Aftab Qamar, Ph.D. Scholar at Institute of Food Science & Nutrition (IFS&N), Bahauddin Zakariya University (BZU), Multan, Pakistan. This research article is a part of the Ph.D. Thesis of Mr. Aftab Qamar. The research work was carried out under the supervision of Dr. Saeed Akhtar Director IFS&N-BZU, Multan. Authors further thank dairy farm owners and other individuals working on dairy farms for their participation and support in this study.

